# Inosine incorporation in DNA nanostructures and 3D DNA crystals

**DOI:** 10.64898/2026.06.06.730639

**Authors:** Arun Richard Chandrasekaran

## Abstract

DNA nanotechnology is based on programmable base pairing, resulting in the precise construction of nanoscale structures. Sequence variability in DNA nanostructure self-assembly is achieved by the use of xeno nucleic acids, chemically modified bases and base analogs. The naturally occurring base inosine, while well studied in RNA editing, has not been used in the context of DNA nanotechnology. In this work, I demonstrate the use of inosine in DNA nanostructures, specifically by incorporating inosine within the duplex regions or junctions of a double crossover DNA motif. In strand displacement and competition assays, I show that canonical complements do not displace inosine containing strands post-assembly but dominate in product formation when competing with inosine containing strands during assembly. Finally, sticky ends with inosine base pairs enable the formation of rationally designed 3D DNA crystals based on the tensegrity triangle motif. Overall, this work shows that inosine is a useful addition to the library of sequence variations in DNA nanotechnology.

---

Beyond the traditional Watson-Crick-Franklin base pairing, the use of non-canonical base pairing, modified bases and DNA analogs has also been demonstrated in DNA nanotechnology.^1–3^ Non-canonical nucleic acids with natural or synthetic structural modifications exhibit enhanced stability and unique functional potential.^4,5^ For example, the Wobble base pair G:U is used to mitigate secondary structures in some nanostructures^6^ and the base pair A:C is used in creating RNA nanostructures with only three bases (A, C, G).^7^ In addition, metal-mediated base pairing has also been used to strengthen mismatches within DNA nanostructures, such as C-Ag-C^8^ and T-Hg-T.^9^ Unnatural base pairs such as 5-Me-isoC/isoG and A/2-thioT have also been incorporated into 6-arm DNA junctions and 6-helix nanotubes, resulting in enhanced thermal stability and resistance to T7 Exonuclease digestion.^10^ Xeno nucleic acids such as threose nucleic acid,^11^ acyclic threoninol nucleic acid,^12^ glycerol nucleic acid,^13^ and peptide nucleic acid^14^ have also been used in DNA-based construction. Recently, newer DNA base analogs have also been created and their use demonstrated in DNA nanotechnology (eg: Z:P).^15,16^ Further, other purine and pyrimidine analogs in addition to the Z:P (such as B:P and D:X pairs of purine analog pairs and S:Z and T:K pyrimidine analog pairs) have been successfully used in several extended tile-based 2D arrays.^17^ The incorporation of such unnatural base pairs in DNA nanostructures provides additional sequence library as well as varied stability and functionality with enzymes such as polymerases and nucleases.^18^

However, a common base, inosine, has not been incorporated into DNA nanostructures and its effect on the self-assembly and other properties of such structures has not been explored systematically. Inosine arises by the deamination of adenosine where an amine group is replaced by a carbonyl (**Fig. 1a**). Inosine behaves like guanosine during base pairing, preferentially forming hydrogen bonds with cytidine. I:C base pairs have reduced stacking energy and form two hydrogen-bond interactions, rather than three in G·C base pairs. Further, poly I:C has been shown to form B-DNA similar to the right-handed B-DNA form of the canonical poly A:T or poly G:C stretches.^19^ Moreover, inosine does not destabilize short DNA-helices when it pairs with either C, U or A,^20,21^ with the stabilities of inosine base pairs being G·C > I·C > I·A > I·G ≈ I·T (**Fig. 1b**).^22,23^ In RNA, A-to-I editing introduces an A-to-G substitution in the RNA sequence without altering the genomic sequence. The broad base-pairing potential of inosine is also naturally observed in cytosolic tRNAs where an inosine is located at the 5′ position (wobble position) of the anticodon,^24^ pairing with C, A or U as originally proposed by Crick.^25^ This base pairing potential, structural similarity to B-DNA when paired and the varied functionality of inosine makes it highly suitable for use in nucleic acid nanotechnology.

**Figure 1.**
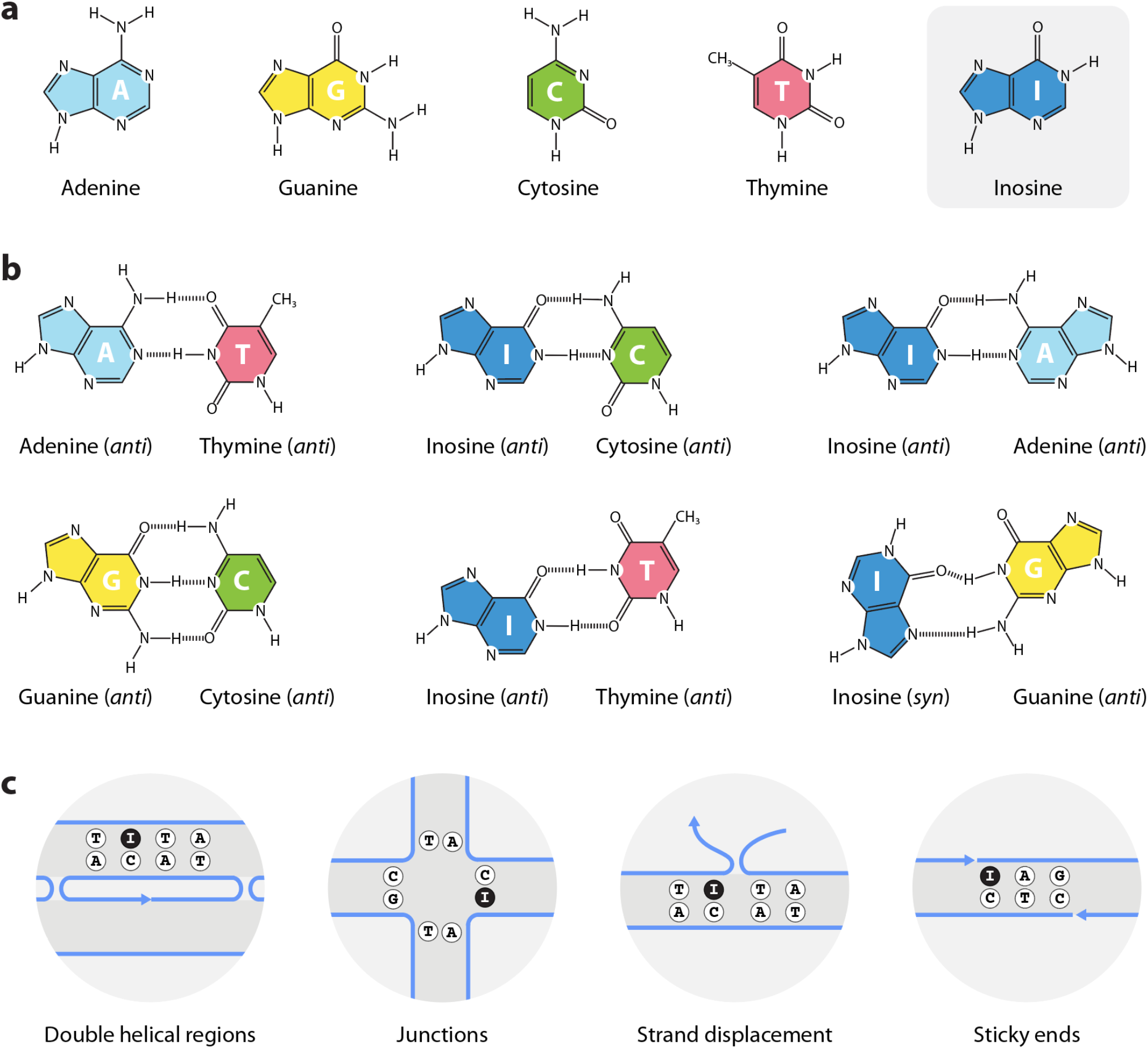
Inosine base pairs. (a) Chemical structures of DNA bases adenine (A), guanine (G), cytosine (C), thymine (T) and inosine (I). (b) Canonical A:T and G:C base pairing (left) and inosine base pairs (right). (c) This study explores inosine incorporation within double helical edges and junctions of DNA nanostructures, effect of inosines in strand displacement and inosine incorporation in sticky ends for hierarchical self-assembly.

Here, I study the effect of inosine incorporation in DNA nanostructures using a model DNA motif – the double crossover (DX) motif. Specifically, inosines were incorporated in the duplex region between crossovers in the structure or at the junction points where the two crossovers occur. In addition, I also study the competition between inosine containing strands and strands containing perfect base pair matches in toehold-less strand displacement. Using the tensegrity triangle motif, I also demonstrate efflcient sticky end cohesion with inosine base pairs in the hierarchical self-assembly of rationally designed 3D DNA crystals.

To study the effect of inosine incorporation in DNA nanostructures, I first used the DX DNA motif,^26^ testing incorporation in two structural regions: one of the double helical regions between the crossovers and within the two junctions. The DX motif has 21 bp between the crossovers, allowing testing of modifications in a 2-turn double helical region constrained by a junction at either end (**Fig. 2a** and **Fig. S1**). I modified the contiguous strand to contain 1, 2, 3 or 4 inosines, with the inosines separated by two bases between them. The DX structures containing different number of inosines can thus be assembled by changing only one of the five component strands. Non-denaturing polyacrylamide gel electrophoresis showed proper assembly of the structures in 1× TAE with 12.5 mM Mg^2+^, with ∼10% decrease in assembly yield for the structure with 4 inosines compared to the control structure without any inosines (**Fig. 2b** and **Fig. S2**). Thermal melting analysis reflected these results, with difference in melting temperature of ∼2-3 °C between the control structure and the structure containing four inosines (**Fig. 2c**). The T_m_ of the structures with 2 or 3 inosines were similar, possibly due to the specific base pairs involved in the inosine positions. Previous studies showed that the effect of inosine incorporation on reverse transcription has also been minimal, possibly due to the tolerance of inosine base pairs in RNA as in DNA observed here.^27^

**Figure 2.**
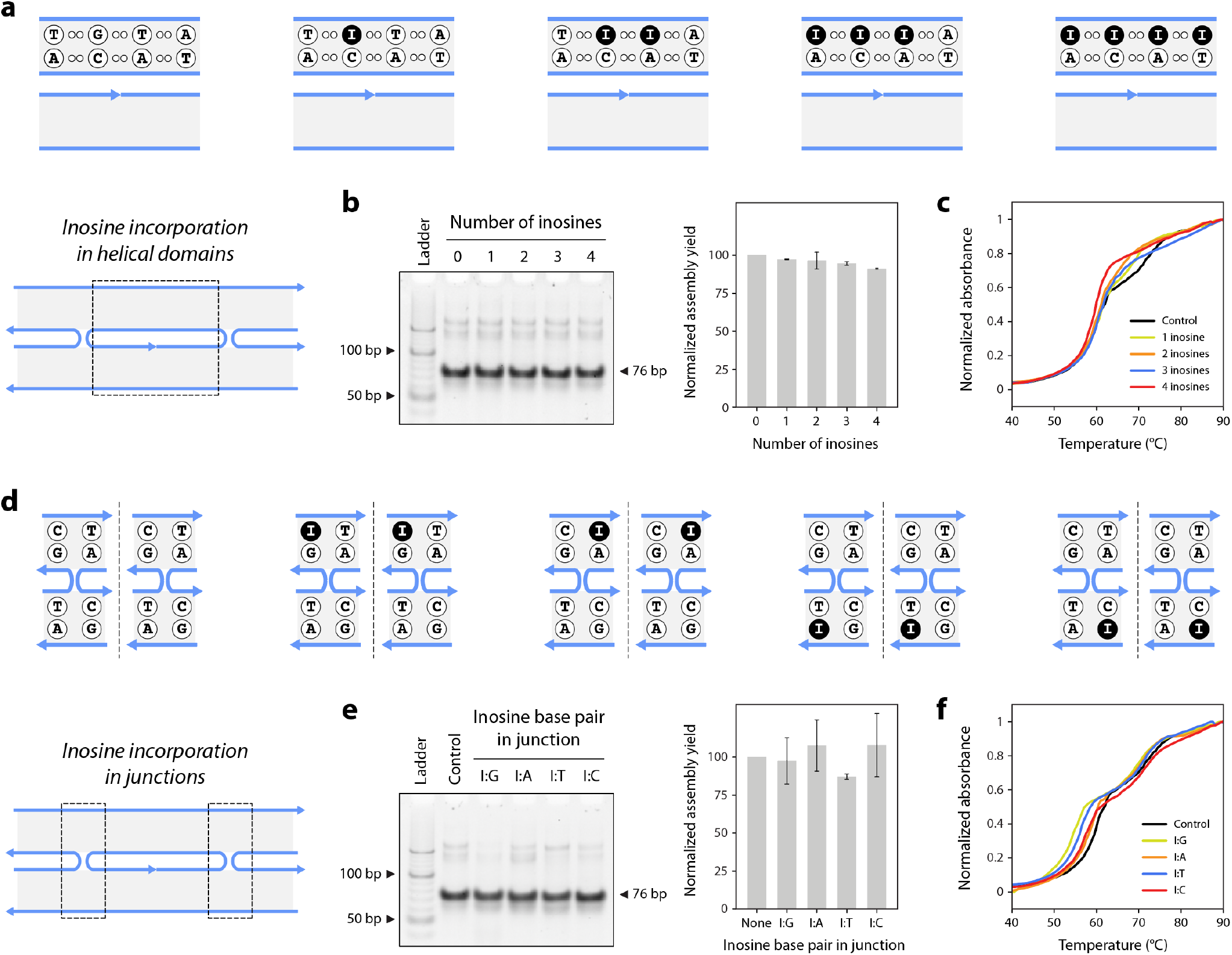
DX DNA motif containing inosines. (a) Schematic showing the number of inosines on one double helical edge of a DX DNA motif. (b) Non-denaturing gel and assembly yields and (c) UV melting curves for DX DNA motif with different number of inosines. (d) Schematic, (e) non-denaturing gel and assembly yield and (f) UV melting curves of DX DNA motif with different inosine base pairs at the two junctions.

Next, I studied the effect of inosines in the junctions of the DX motif. The sequence of the DX at the junction allows each type of base in the core four positions to be replaced by an inosine, i.e. I:G instead of C:G, I:T instead of A:T, I:C instead of G:C and I:A instead of T:A (**Fig. 2d** and **Fig. S3**). This design allows the study of inosine incorporation within a DNA junction, while also comparing the different effects on the four types of base pairs. Compared to the control structure without inosines, the assembly and T_m_ of the inosine-containing structures were similar, indicating minimal effect when only one of the base pairs of the junction was modified (**Fig. 2e-f**).

I then studied the competition between strands containing the canonical base pairs and inosine base pairs. In one of our recent works,^28^ we demonstrated mismatch induced strand displacement where the instability in a duplex due to mismatches allowed strand displacement to occur without the presence of a toehold. I tested a similar effect here to determine whether the inosine base pairs acted as mismatches or were less stable compared to the canonical base pairs, thereby allowing a full complement to displace it. I designed duplexes formed by strand X* and a complement containing 1, 2, 3 or 4 inosines (X-I_n_). To the assembled duplex, I added an invading strand with a perfect match to observe whether the inosine containing strands are displaced. I included poly T extensions at either end of the invading strand (X_T_) to allow convenient analysis by differential gel migration of the initial and final duplexes (**Fig. 3a** and **Fig. S4**). I observed that there was minimal strand displacement compared to the control structure, indicating that the inosine base pairs did not behave like typical mismatches in the structure and (**Fig. 3b** and **Fig. S5**).

**Figure 3.**
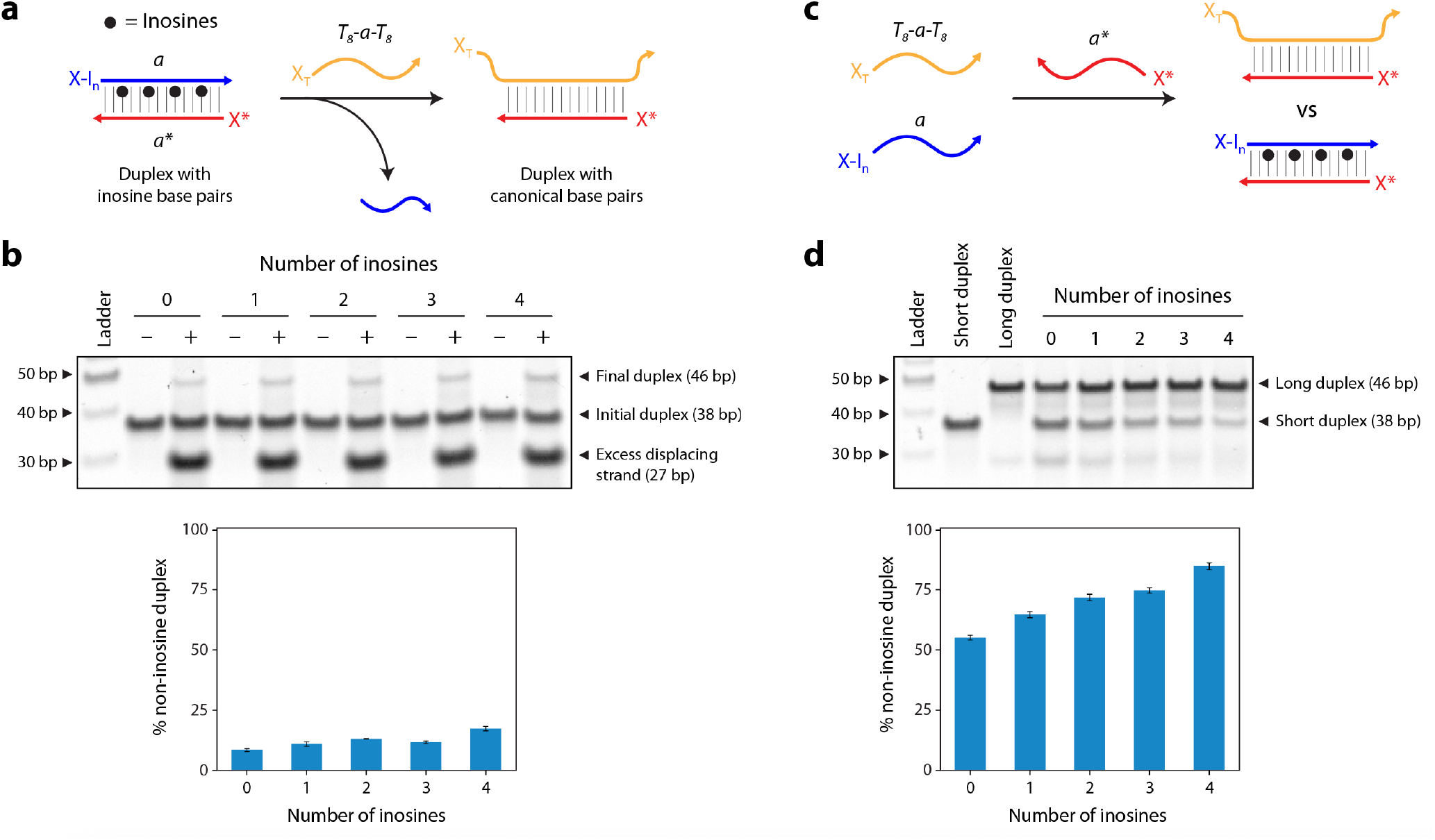
Strand displacement and competition. (a) Displacement of inosine-containing strand from a pre-assembled duplex. The invading strand X_T_ contains a full complement to strand X* and additional Ts at the two termini to differentiate the final complex from the initial duplex on a gel. (b) When inosine containing complement and a full complement compete for the same binding partner (X*), the strand containing canonical bases is preferred. Strand containing higher number of inosines is less preferred.

Based on this result, I then tested whether inosine containing strands and canonical complements would compete in DNA duplex formation (**Fig. 3c**). I mixed component strands with (X-I_n_) and without (X_T_) inosines, both of which are complementary to strand X*, and annealed the solution. Again, the conventional complement had polyTs on the termini to allow gel-based identification of the products. The control without inosines also showed 50% of each product, due to the equilibrium between both complements, a feature we have observed in our earlier work.^28^ Results showed that in each case, the complement with the canonical bases formed the major product, with up to ∼85% yield compared to only ∼15% yield for the duplex containing four inosines (**Fig. 3d** and **Fig. S5**). Notably the duplex product with the inosine containing strand decreased as the number of inosines increased, indicating that when directly competing with a complementary partner, the strand with conventional base pairs is preferred.

Next, I tested the incorporation of inosine in sticky ends that allow the hierarchical assembly of DNA nanostructures. As a model system, I chose 3D DNA crystals self-assembled from the tensegrity triangle DNA motif, a system I’ve extensively worked on and well-studied by other groups as well.^29–42^ The structure contains three double helical edges (each 31 bp in this case) that are connected at the vertices by a four-arm junction (**Fig. 4a**).^43^ The structure is inherently three-dimensional, with an over-and-under arrangement of the helical edges. The edges are further tailed by sticky ends that allow the triangles to connect to each other, resulting in the formation of a three-dimensional (3D) DNA lattice. Typically, the triangles are tailed by 2 nucleotide sticky ends in crystal formation; however, other sticky end lengths and sequences have also been studied.^33^ To analyze the effect of inosine, I modified one of the sticky end bases to inosine (**Fig. S6**). This allowed me to study whether inosine containing sticky ends allow hierarchical assembly of DNA motifs, especially in this 3D arrangement where micrometer sized crystals contain trillions of the starting triangle motif. There is only one occurrence of an inosine containing sticky end in DNA nanostructures, where an II:CC sticky end was used in a similar crystal assembled from 2-turn tensegrity triangles (a study I was part of).^33^ Here, I use tensegrity triangle DNA motifs with 1-nt and 3-nt sticky ends containing inosines.

**Figure 4.**
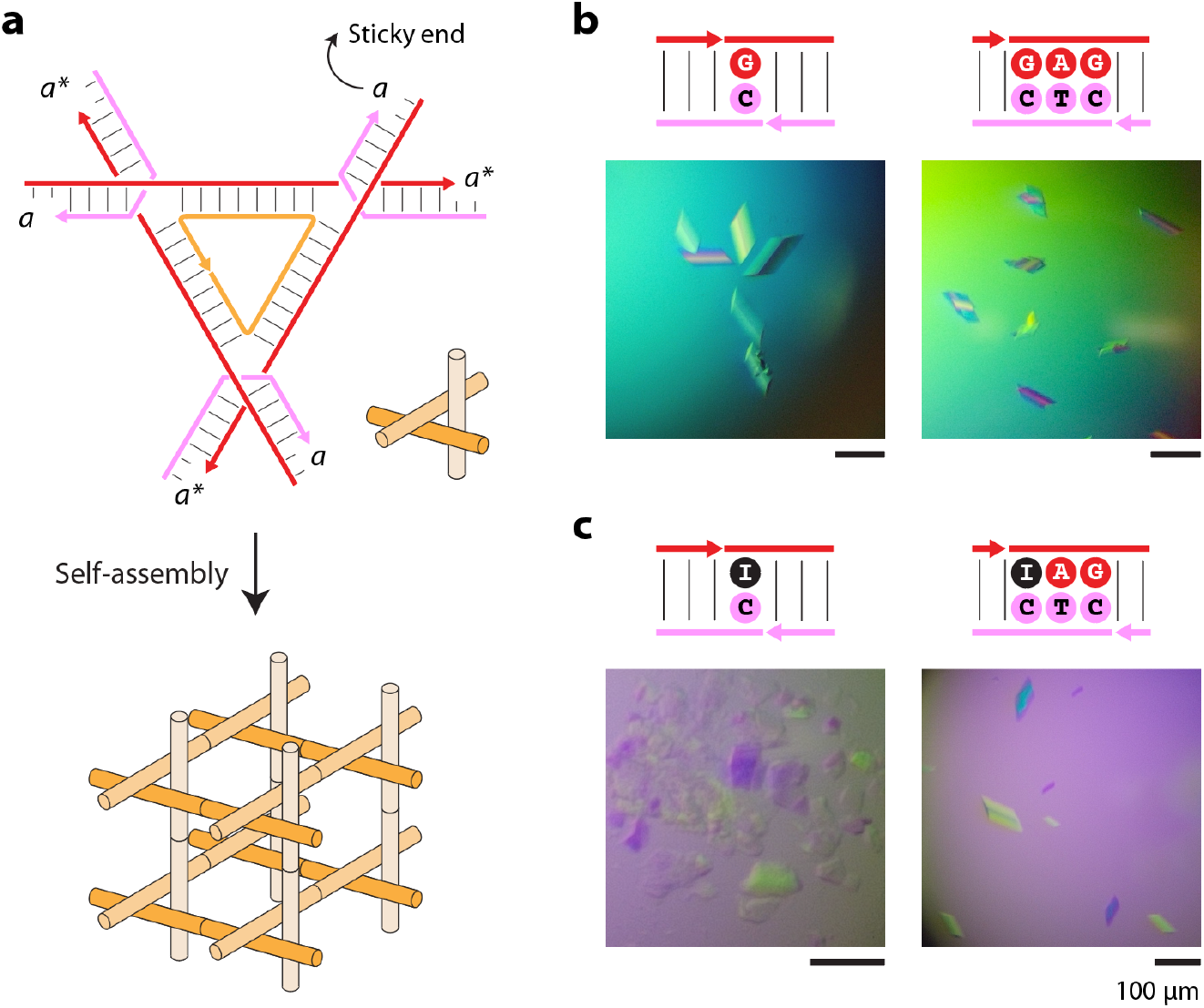
Hierarchical assembly of 3D DNA crystals. (a) Schematic of the tensegrity triangle motif and self-assembly via sticky end cohesion into 3D crystals. Each edge of the motif contains 31 bp regardless of the sticky end length. Crystals obtained from tensegrity triangle motifs using sticky ends without (b) and with inosine (c).

Non-denaturing gels showed proper assembly of each version of the triangle, with a single band corresponding to the structure (**Fig. S7**). For the structure with 3-nt GAG:CTC sticky end, I also observed additional DNA in the wells, a result that corresponds to stronger sticky end cohesion resulting in hierarchical structures. I set up crystals using the triangles with different sticky end lengths, both with and without inosines. I observed crystal growth in all the triangular assemblies, indicating that the sticky ends allowed self-assembly of the triangles into crystals (**Fig. 4b**). Inosine containing sticky ends also allowed crystal formation, with the crystal habit observed resembling those of the controls and the typical shape observed for these designer DNA crystals in our previous studies (**Fig. 4c**). In addition to the one prior report of an II:CC sticky end, I show in this work that even with a single nucleotide pair, where the interaction is I:C, the triangles are still able to self-assemble into 3D crystals.

Overall, this work shows that inosine is a useful addition to the library of sequence variations in DNA nanotechnology. While other unnatural base pairs with the anti-syn glycosidic conformation have been developed, inosine is a natural base that prefers the syn confirmation when pairing with guanine.^44^ With increasing efforts on developing nanostructure based mRNA delivery,^45,46^ introduction of inosine may be useful in triggering specific cellular responses.^47^ Inosine-induced base pairing diversity in RNA is well studied but more studies are needed to understand the structural effects of inosine base pairs in DNA helices. The effect of G to I substitution in A-RNA and B-DNA have been shown to depend on the different topologies of these helices, with the different electrostatic interactions between the base pairs and the negatively charged backbone playing a role.^48^ Increasing use of inosines in DNA related applications, such as a the work reported here, may initiate more detailed studies on the topic.

## Supporting information

Experimental procedures, additional results and DNA sequences used.

## Acknowledgments

Research reported in this publication was supported by the National Institutes of Health (NIH) through National Institute of General Medical Sciences (NIGMS) under award number R35GM150672 to A.R.C. This manuscript is the result of funding in whole or in part by the National Institutes of Health (NIH). It is subject to the NIH Public Access Policy. Through acceptance of this federal funding, NIH has been given a right to make this manuscript publicly available in PubMed Central upon the Official Date of Publication, as defined by NIH.

