## Supplementary material for "Inosine incorporation in DNA nanostructures and 3D DNA crystals": SI

### Methods

***Preparation of DNA complexes.*** DNA strands were purchased from Integrated DNA Technologies (IDT). DNA complexes were annealed in TAE buffer containing 40 mM tris base (pH 8.0), 20 mM acetic acid, 2 mM EDTA and 12.5 mM magnesium acetate (concentrations in 1× buffer). DX samples were annealed in a thermal cycler with the following steps: 90 °C for 3 minutes, 65 °C for 20 minutes, 45 °C for 20 minutes, 37 °C for 30 minutes, 20 °C for 30 minutes, and the solution was then cooled to 4 °C. Duplexes were annealed from 90 °C to 20 °C over ~30 min. Tensegrity triangle nanostructures were annealed from 90 °C to 20 °C over 2 days.

***Gel electrophoresis.*** Non-denaturing polyacrylamide gels were prepared using 19:1 acrylamide to bisacrylamide stabilized solution (National Diagnostics) in 1× TAE-$Mg^{2+}$ buffer. DNA samples were mixed with loading dye containing bromophenol blue and glycerol before loading into gels. Gels were run at 4 °C in 1× TAE-$Mg^{2+}$ running buffer and stained with 0.5× GelRed (Biotium) and destained in water. Gels were imaged on a Bio-Rad Gel Doc XR+ imager using the default settings for GelRed with UV illumination and analyzed using ImageLab software (Bio-Rad). The assembly yield was quantified as the fraction of the intensity of the band corresponding to the structure compared to the total intensity of all the bands in the lane.

***Thermal melting studies.*** UV thermal melting experiments were performed in a Cary 3500 UV-Visible Spectrophotometer equipped with a temperature controller. Melting curves were acquired at 260 nm by heating and cooling from 20 °C to 90 °C at a rate of 0.5 °C/min. The data were fitted to the Boltzmann function using OriginPro. Melting temperatures were determined from the first derivative of the fitted melting curves.

***Crystallization.*** 5-6 μl of annealed tensegrity triangle structures were equilibrated against a 600 μl reservoir of 1.75 M ammonium sulfate in hanging drops set up at 20 °C. Crystals were also set-up using the DNA solution in hanging drops and the entire set up placed in a thermal gradient from 60 °C to 20 °C. Crystals were imaged using a Zeiss SteREO Discovery V12 microscope.

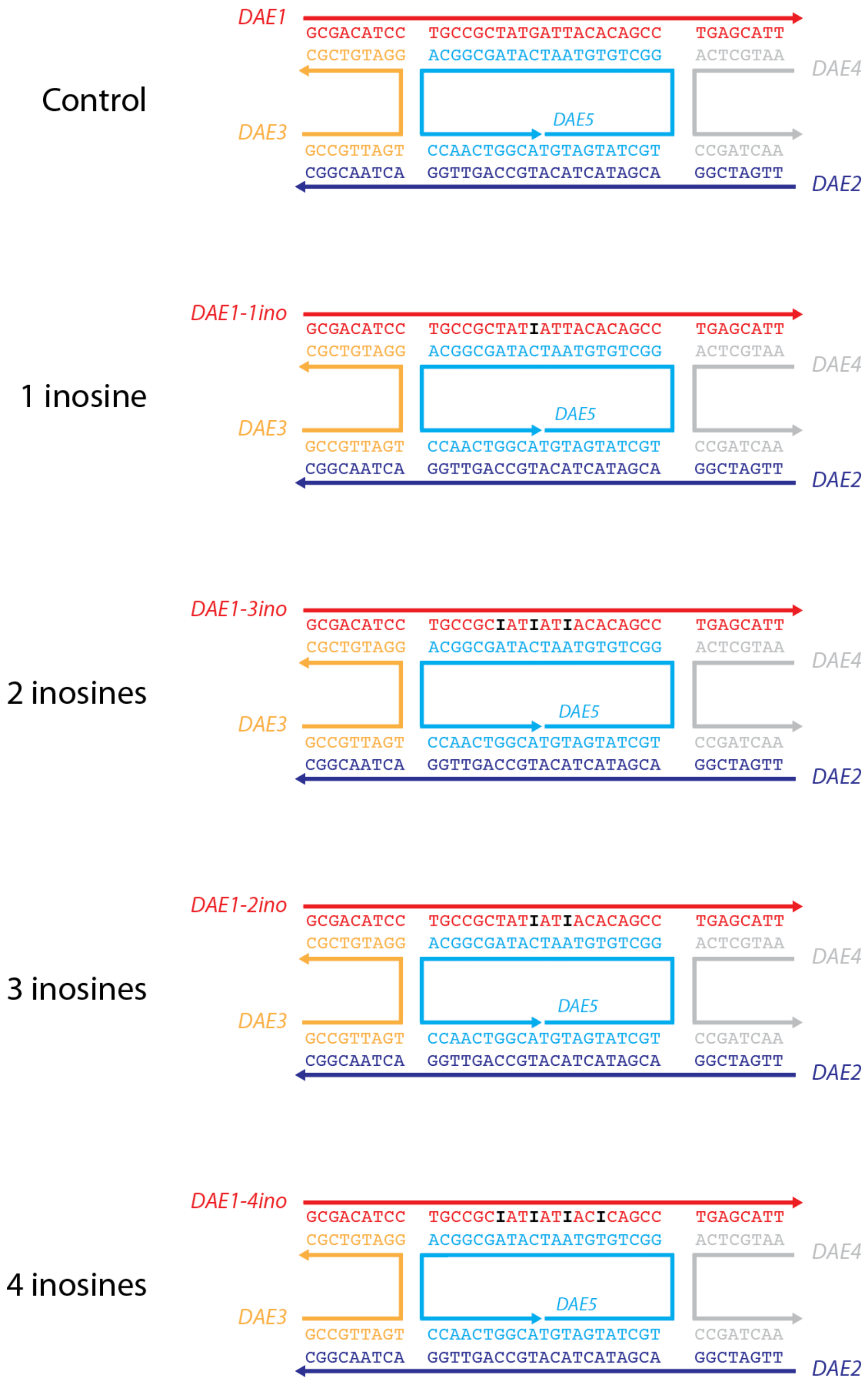

**Figure S1**. Sequences used in the DX motif. Strand 1 was modified to contain 1-4 inosines.

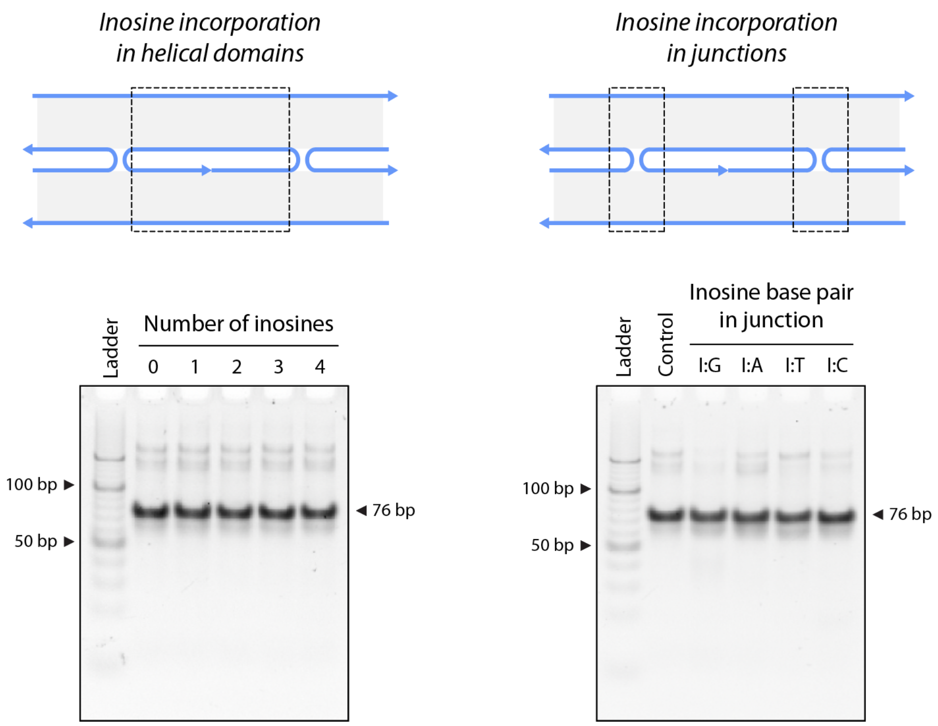

**Figure S2**. Assembly of DX DNA motif containing inosines. Full images of gels shown in Figure 2.

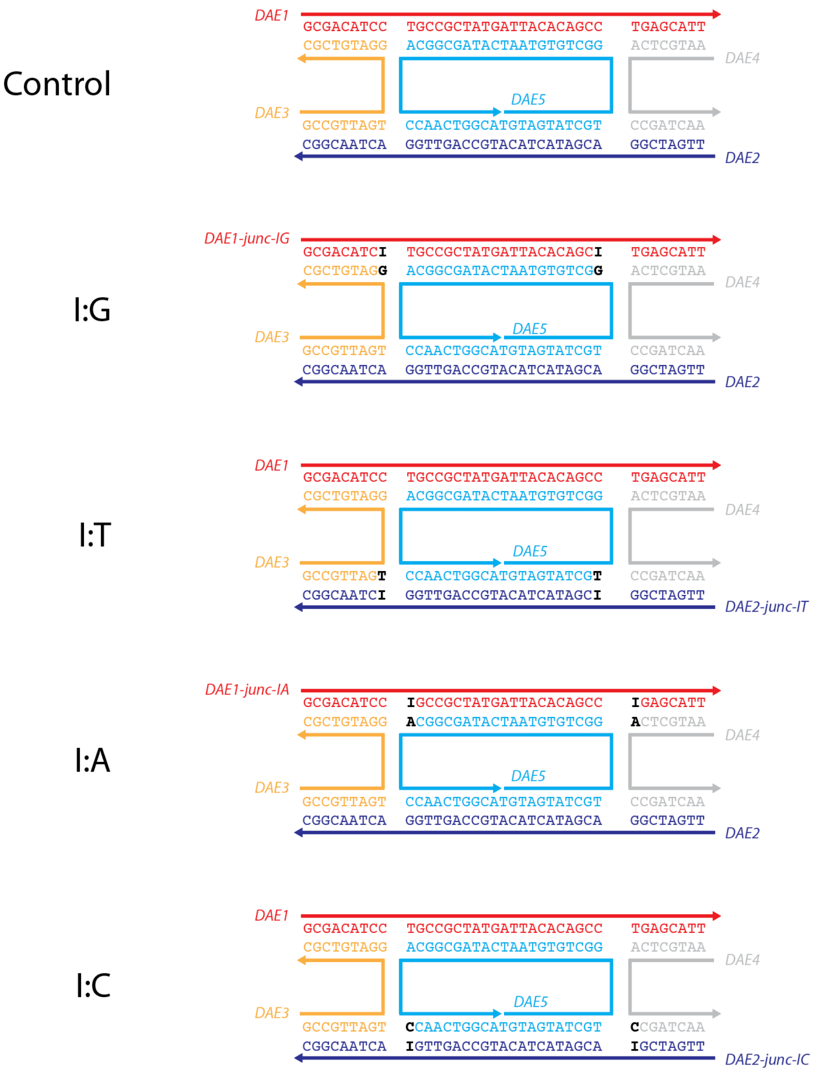

**Figure S3**. Sequences used in the DX motif, with strands modified to incorporate inosine in the junctions.

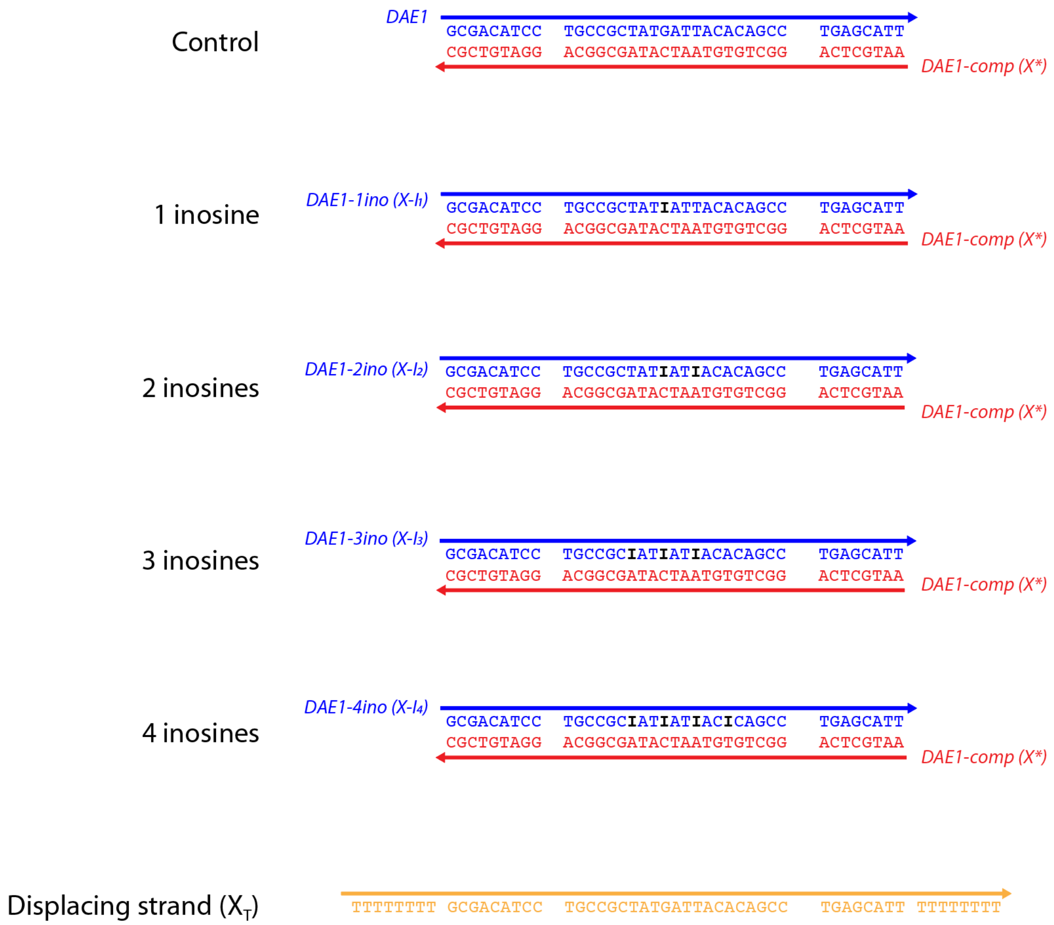

**Figure S4**. Sequences used in the strand displacement experiments. The displacing strand $X_T$ contained additional Ts at the termini to allow convenient readout by gel electrophoresis.

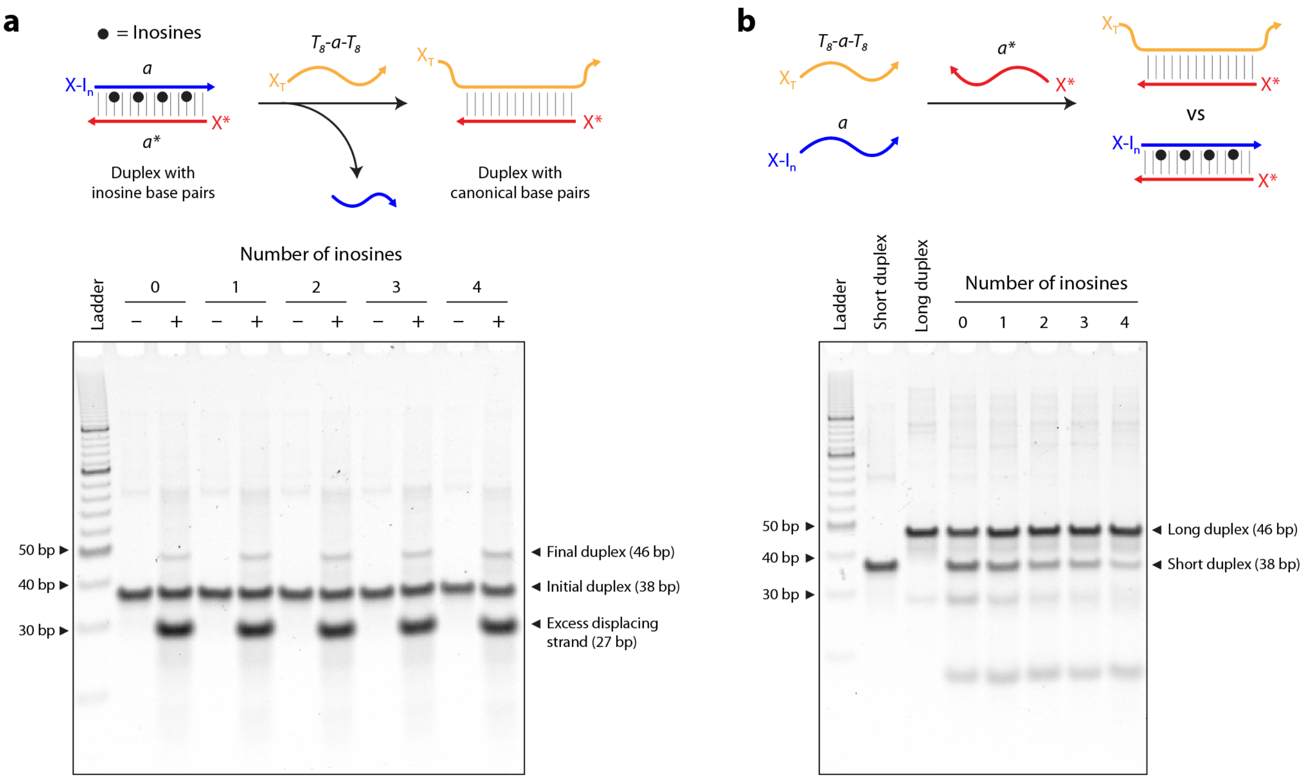

**Figure S5**. Strand displacement (a) and competition (b) in duplexes containing inosines. Full images of gels shown in Figure 3.

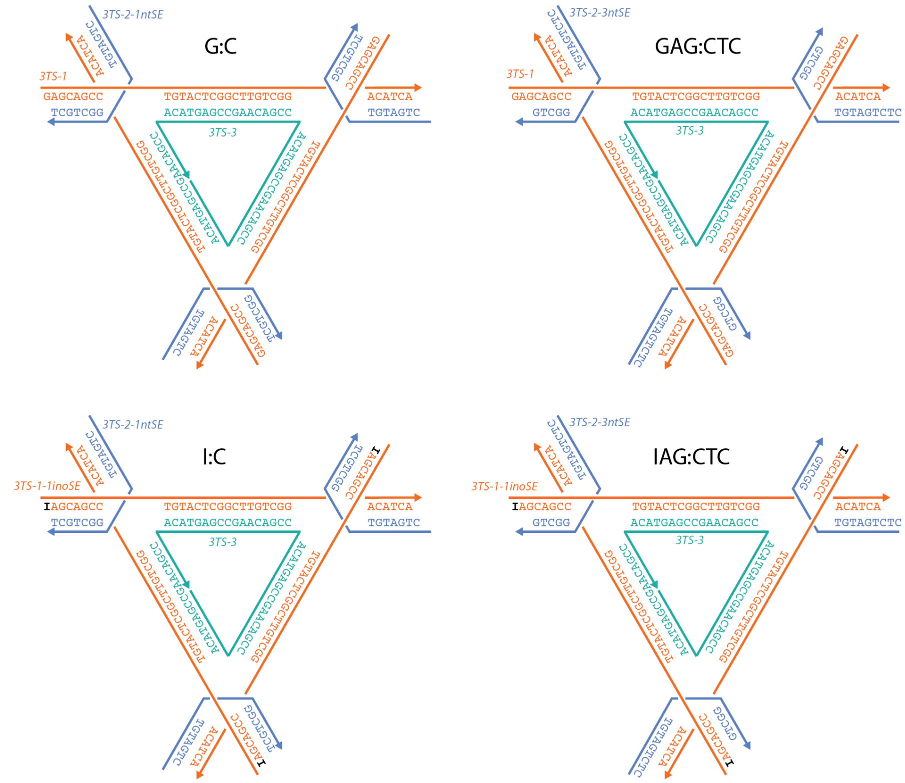

**Figure S6**. Sequences used in the tensegrity triangle motif. Strand 1 was modified to contain an inosine and strand 2 was modified to obtain different sticky end lengths. The overall length of the duplexes in each edge was constant at 31 bp in all variations.

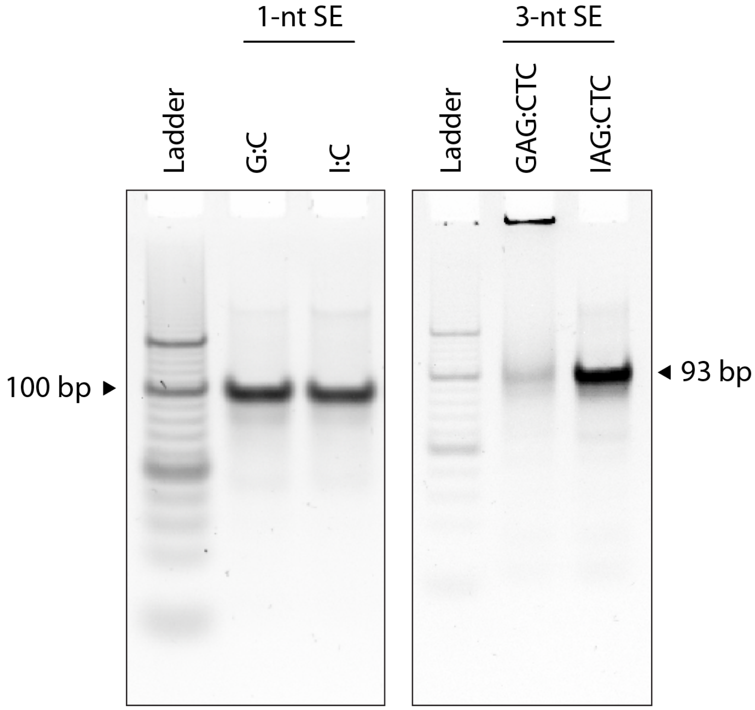

**Figure S7**. Assembly of the tensegrity triangle DNA motif containing inosines in the sticky ends.

| **Double crossover (DX) motif** | |
|---|---|
| DAE-1 | GCGACATCCTGCCGCTATGATTACACAGCCTGAGCATT |
| DAE-2 | TTGATCGGACGATACTACATGCCAGTTGGACTAACGGC |
| DAE-3 | GCCGTTAGTGGATGTCGC |
| DAE-4 | AATGCTCACCGATCAA |
| DAE-5 | TGTAGTATCGTGGCTGTGTAATCATAGCGGCACCAACTGGCA |
| DAE1-1ino | GCGACATCCTGCCGCTAT**I**ATTACACAGCCTGAGCATT |
| DAE1-2ino | GCGACATCCTGCCGCTAT**I**AT**I**ACACAGCCTGAGCATT |
| DAE1-3ino | GCGACATCCTGCCGC**I**AT**I**AT**I**ACACAGCCTGAGCATT |
| DAE1-4ino | GCGACATCCTGCCGC**I**AT**I**AT**I**AC**I**CAGCCTGAGCATT |
| DAE1-junc-IG | GCGACATC**I**TGCCGCTATGATTACACAGC**I**TGAGCATT |
| DAE1-junc-IA | GCGACATCC**I**GCCGCTATGATTACACAGCC**I**GAGCATT |
| DAE2-junc-IT | TTGATCGG**I**CGATACTACATGCCAGTTGG**I**CTAACGGC |
| DAE2-junc-IC | TTGATCG**I**ACGATACTACATGCCAGTTG**I**ACTAACGGC |
| **Duplexes** (used with DAE1 with and without inosines) | |
| DAE1-comp (X*) | AATGCTCAGGCTGTGTAATCATAGCGGCAGGATGTCGC |
| DAE1-8T ($X_T$) | TTTTTTTTGCGACATCCTGCCGCTATGATTACACAGCCTGAGCATTTTTTTTTT |
| **Tensegrity triangle motif** | |
| 3TS1 | GAGCAGCCTGTACTCGGCTTGTCGGACATCA |
| 3TS1-1ino | **I**AGCAGCCTGTACTCGGCTTGTCGGACATCA |
| 3TS2-1nt SE | CTGATGTGGCTGCT |
| 3TS2-2nt SE | TCTGATGTGGCTGC |
| 3TS2-3nt SE | CTCTGATGTGGCTG |
| 3TS3 | CCGAGTACACCGACAAGCCGAGTACACCGACAAGCCGAGTACACCGACAAG |

**Table S1**. Sequences used in this work (written 5′-3′).
